# Putting on academic armor: How Black physicians and trainees take stances to make racism visible amidst publishing constraints

**DOI:** 10.1101/2022.12.27.521686

**Authors:** M. Johnson, L.A. Maggio, A. Konopasky

## Abstract

**Introduction:** While there are a growing number of empirical studies of Black physicians’ and trainees’ experiences of racism, there are still few accounts from a first-person perspective. These personal commentaries or editorials require taking a delicate stance, balancing the professional, social, and individual. Black authors in the medical publishing space, who already experience microaggressions and racial trauma in their work spaces, must put on their academic armor to further experience them in publishing spaces. Medicine and medical education interpellate each of us, including the authors and readers of this piece, as a particular kind of subject. Beginning with a narrative grounding in the authors’ personal stances, this study seeks to understand the stances Black physicians and trainees take as they share their personal experiences of racism while protecting the institution of medicine from racism’s irreparable harms.

**Methods:** The authors searched four databases identifying 29 articles authored by Black physicians and trainees describing their experiences. During initial analysis, the authors identified three sets of discursive strategies: *identification* (introducing and tracking themselves throughout the article); *intertextuality* (dialoguing and drawing on other “texts” beyond the focus of the articles); and *space-time* (social constructions of space and time linked to a network of social practices). The authors used a Google form to capture these strategies and direct quotes from the articles. Throughout the study, the authors reflected on their *own* stances in relation to the experience of conducting the study and its findings.

**Results:** Authors engaged in stance taking, which aligned with the concept of donning academic armor, by evaluating and positioning themselves with respect to racism and the norms of academic discourse in response to ongoing conversations both within medicine and in the broader US culture. They did this by (a) positioning themselves as being Black and, therefore, qualified to notice and name personal racist experiences while also aligning themselves with the reader through shared professional experiences and goals, (b) intertextual connections to other related events, people, and institutions that they–and their readers–value and (c) aligning themselves with a hoped-for future rather than a racist present.

**Discussion:** Because the discourses of medicine and medical publishing interpellate Black authors as Others they must carefully consider the stances they take, particularly when naming racism. The academic armor they put on must not only be able to defend them from attack, but help them slip unseen through institutional bodies replete with mechanisms to eject them. In addition to analyzing their own personal stance, authors leave readers with thought provoking questions regarding this armor as they return to narrative grounding.

## Narrative Grounding: Starting with Reflexivity

We begin this piece about the stances Black physicians and trainees take in their published writing by attempting to be transparent about our own stances. We voice here our own understandings about how medicine and medical publishing *interpellate* (i.e., constitute) us as certain kinds of subjects.^1^ We offer these explications of the different academic armor we each put on as we enter this space of publishing to help readers understand our unique stances and how they shape this work.

### Author 1: Reading Between the Lines

As a Black woman medical student at a predominantly white institution (PWI), I experienced and continue to experience racism in several aspects of my life. Academic publishing has exposed me to a world of scientific expression, where using one’s experience to teach a community is non-traditional and usually not enough on its own to be accepted as “data. ” As such, I view research with personal experiences woven in through a lens of reading between the lines. I assume that what is printed is carefully crafted and that there is almost always more to the story. This is second nature as I tend to identify with the majority of experiences shared by Black physicians/trainees and I am familiar with how to frame that narrative. My lens and my curiosity regarding how other Black physicians and trainees experience and display that experience in the literature are what started this collaboration.

### Author 2: Seeing Scholarly Publishing as Values-Laden

As a white woman full professor and deputy editor-in-chief of a journal, I have over a decade of experience as an author, mentor, and gatekeeper in a tradition of scholarly communication in which an article looks a certain way (i.e., highly structured and generally impersonal). Until recently, I was ignorant of how I have contributed to and perpetuated this system. Simply, I have failed to recognize scholarly publishing as a values-laden system, which privileges those that share a background and approach mirroring my own. The opportunity to publish Authors 1 and 3’s commentary^2^ in which they share their personal reflection on publishing, raised my awareness of scholarly publishing’s norms and motivated me to collaborate on this research.

### Author 3: Language as Clue to Racial Strictures

As a white, heterosexual, cisgendered associate professor with few Black acquaintances (let alone friends) and numerous opportunities to share my voice through publishing, initially I had purely book-based knowledge of racism in academic publishing. Mentoring Author 1 has allowed me to use academic spaces to bear witness to her experiences of racism, disrupting these spaces along with her. Yet I continue to commit racist microaggressions against her^2^, so I see Author 1’s lens as having primacy in this work. Moreover, as a linguist, I believe in the power of language to oppress and to free, so I approach research through the lens of language. In particular, my interest here has been in what the linguistic choices of Black physicians/trainees who publish might say about the racial strictures–explicit and implicit–they are writing under.

There is an inextricable link between Black physicians’ professional identity and their social identity. Yet first-hand experiences from this group are rarely published in academic journals.^3^ It is more “acceptable” for those experiences to be portrayed through a third-person lens in alignment with the expectations of structured academic discourse^4^ than a first-person voice.^3^ As a result, much of the available literature about Black physicians’ and trainees’ experiences are published as empirical studies. We read *about* Black physicians and trainees, from a scientific distance, but we do not often listen directly *to* them. To do so might threaten the hopeful future we build in our academic publishing, a future of findings to be implemented and studies to come. To do so, would be to give space for complaints of racism, for *testimony to* racism.^5^ Yet this direct testimony is vital; it reveals the experience of being a Black body in a white institution, it “provides a *phenomenology of the institution*”.^5(p.41)^ Unfortunately, however, this testimony is constrained: to protect the institution, Black voices are expected to obscure these harms and even project happiness at being given a place there.^5,6^ In order to truly understand the devastating experiences of racism for Black physicians and trainees and the role of the institution in obscuring them, we must listen to their testimony and understand exactly *how* they make their voices heard.

Results from empirical studies provide some explanation, albeit incomplete, for the lack of first-hand accounts from Black physicians and trainees. Studies have demonstrated the negative experiences of Black physicians, including the psychological and physical effects of ongoing microaggressions^7^ and a “minority tax”, which places extra burdens on them based on their racial identity.^8^ This also suggests a lack of psychological safety in their training and work environments because there are so few physicians who share and understand their history.^3,8^ Their negative experiences are also partially due to racial trauma as a result of witnessing and experiencing racial violence. Racial violence can include both direct acts of racism, such as hate crimes or workplace discrimination, as well as more systemic forms such as health disparities and financial inequity.^3,9,10^ This lack of safety, among other factors, can lead to the cumulative effect of racism on an individual’s mental and physical health and racial trauma, which we know can lead to hypervigilance, withdrawal and emotional exhaustion.^11^ While a plethora of research has cataloged racial violence in healthcare as a whole, there needs to be a space held for those experiencing this systemic racism to share their lived experiences and have them accepted as valid contributions to academia.

The nature of academic discourse is another reason for the lack of first-hand accounts. Publishing a journal article is considered the primary communication mechanism in academia. Beginning in the 1600s, journal articles were freeform in presentation with limited guidelines for article structure until the 1940s.^12^ At that time, the IMRaD format was introduced, which instructed authors to structure articles in the following sections: introduction, methods, results and discussion. The IMRaD format is the dominant format of journal articles,^4^ which is unsurprising since there is a risk of rejection when straying from the structure.^13^ The IMRaD format puts authors under strict constraints, including not only how they organize their articles, but also how they develop their argumentation and select the words to represent their own voice.^14^ In medicine, this has meant that “the practitioner-author has gradually been replaced with the academician-author,” no longer sharing personal experience in medicine, but scientific contributions.^15(p.317)^ In other words, medical publishing has moved from testimony to science.

Researchers have identified a tension between the expectations of the academic community that authors will follow certain rules and expectations (i.e. adherence to IMRaD and a “scientific” format) and the desire by authors to “claim their agency and develop their uniqueness”^16(p.83)^ Publishing is a critical national and international space where physicians and trainees negotiate their professional identity where they are challenged to hew to conventions and use their voice to construct a particular social identity.^16^ This negotiation is particularly difficult for Black authors for two reasons: first, they have no choice in their interpellation as Black–and therefore, Other–so they must address that imposed social identity. Second, naming racism breaks the convention of protecting the institution while *not* naming it and projecting happiness is to obscure your social identity as Black in medicine.^5,6^ Thus, Black authors must carefully take *stances*, positioning themselves simultaneously vis-à-vis racism and the institutions of medicine and the academy.^17^ The purpose of this study is to explore the stances Black physicians and trainees take as they negotiate sharing their personal experiences of racism while protecting the institution of academic medicine from racism’s irreparable harms.

## Methods

We conducted a qualitative analysis of journal articles published by Black physicians and trainees in medicine about their personal experiences of racism. The aim was to examine the 17 various *stances*–public acts of evaluation in order to position the self with respect to others^17^ – these authors took in order to negotiate the academic and medical conventions around harming institutions by referencing racism.^5^ As this study focuses on published literature and does not include human subjects, we did not submit for ethical review.

To identify articles, a medical librarian systematically searched PubMed, CINAHL, PsycINFO and Web of Science using a combination of keywords tailored for each database. Search terms included, but were not limited to, terms related to race (e.g., racial, racism, Black, African American) AND medicine (e.g., medical, healthcare). (See Appendix A for complete search terms). We limited the search to articles published between 2018-2021. We selected this start date to coincide with the period before the murder of George Floyd (as well as other Black individuals: Brionna Taylor, Ahmed Arbery) due to the level of increased public outcry and the public push for an increase for “diversity” in multiple arenas around that time. Searches were conducted on November 9, 2021. All results were transferred to Covidence, a review software, and duplicates removed.

Two authors independently screened all article titles and abstracts for inclusion. Articles were included if at least one author identified in the article as a Black physician or medical trainee and narrated their experience as a Black individual in medicine using first person (“I” or “we”) at least once. Empirical articles were excluded (e.g., a survey study about Black medical students). For those articles that passed title/abstract screening, we reviewed the full-text in duplicate to confirm inclusion.

### Analysis

We began by reading the same three articles, contextualizing them within the literature and our own experiences, and then we discussed the strategies we saw authors using to take stances that allowed them to name and share personal experiences of racism. We identified three sets of discursive strategies authors used: *identification* (introducing and tracking themselves throughout the article);^18^ *intertextuality* (dialoguing and drawing on other “texts” beyond the focus of the articles);^19^ and *space-time* (using social constructions of space and time linked to a network of social practices).^19^ Each of these strategies were ways authors engaged in stance taking–positioning and aligning themselves vis-a-vis other people, entities, and events.^17^ We created a Google form with data collection fields for each of these three discursive strategies and direct quotes demonstrating them. Author 1 coded all articles and Authors 2 and 3 each coded half. The team met several times via Zoom to interpret our emerging findings and discuss our *own* stances towards this work (see narrative grounding above).

### Limitations and Considerations

While we conducted a broad search of the literature, the authors of the included studies are primarily based at institutions in the U.S., which may indicate that our findings do not speak for non-US authors. Additionally, we were able to analyze only those accounts that were published. It is likely that for the 29 articles analyzed that there are scores more that were not accepted for publication. Moreover, as published articles, we were unable to determine the effect of copyediting and peer review, which may have influenced the way in which ideas were presented. Last, but not least, in this article we attempt to be highly transparent about the backgrounds and views of our author team, however, a different author team may have approached this from a different angle.

## Results

We analyzed 29 articles (See Appendix B for a complete listing of all articles), which were published in a variety of journals (n=24) with the *Academic Medicine* journal publishing the most (n=5). (See Appendix C for the individual coding of each article). Notably nearly all journals only published a single account (n=21). The majority of articles were described as commentaries, viewpoints, perspectives, editorials, and letters. In terms of presentation, many did not include an abstract and for those that did the abstract was unstructured. In terms of length, articles were on average 3.2 (MED=3, St. Dev = 2.4) pages. The majority of articles included cited references. However, in one instance, authors had published a letter to the editor, in which the journal did not permit references for this publication type. However, the authors instead provided a recommended list of “works that could have been cited” some of which were featured in the article.

All articles included at least one author who was either a health professions trainee (e.g., medical student) or physician with co-authors representing a range of individuals, including allied health professionals, educators, historians, and researchers. Authors were located primarily in the United States (n=63). The largest author team consisted of 14 authors, which was an open letter co-signed by members of the Black Feminist Health Science Studies Collective.^20^ Six articles were written in dyads. Several articles were written by author groups (e.g., the Allied Black Students located at a medical school).^21^

In our analysis we identified four stances that Black physicians and trainees take as they negotiate sharing their personal experiences of racism while protecting the institution of academic medicine from racism’s irreparable harms. These stances included: stance as a social act, identification and positioning of self, intertextual connections, and promissory space-time endings.

### Stance as a Social Act

In order to share experiences of racism–a word which, in itself, can seem threatening to an individual, a department, or an institution–participants had to negotiate with readers a shared *stance object*, the issue about which they were taking a stance (i.e., racism^17^). While we included these articles because we interpret racism to be the commonly shared stance object among them, it is rarely mentioned explicitly in the articles. Instead, authors carefully aligned themselves with their (likely predominantly white readers if we consider the demographics of medicine^22^) through, (a) positioning themselves as being Black and, therefore, being qualified to notice and name personal racist experiences while also aligning themselves with the reader through shared professional experiences and goals, (b) intertextual connections to other related events, people, and institutions that they–and their readers–value and (c) aligning themselves with a hoped-for future rather than a racist present.

In presenting our findings below, we emphasize author voices by beginning with them and listing their names, referring back to these quotes as we discuss our interpretations.

### Identification and Positioning of Self

“I have served as Associate Vice Provost for Diversity, as the first Black chair of a clinical department at an Ivy-League university (2014-2018), and lead community engagement and health equity efforts.” -Doubeni^23(p.1387)^

“As a Black physician-scientist of Nigerian heritage, whose doctorally trained parents instilled a love for learning and a passion for service in their 7 daughters (1 lawyer/accountant, 3 subspecialist medical doctors, 2 pharmacists, 1 optometrist/public health specialist) and 3 sons (engineers/information technology specialists), it is saddening to learn that education, professional experience, or social status does not automatically overcome bias in our society.” - Mezu-Nbisi^24(p.4)^

“I can remember the first time that I realized that I was Black and that being Black in America would lead to a life of discrimination. I was in the first grade in urban Alabama, and two of my White classmates were sharing lip balm as we stood in the lunch line. I asked, ‘Can I have some lip balm?’ ‘No, you can’t have any, because you’re Black,’ my classmate replied.” - Griffin^25(p.1044)^

“As a Black woman in medical school, I have had to cultivate ways to protect myself emotionally from the vicarious trauma triggered by witnessing the racially charged murders of Ahmaud Arbery, Breonna Taylor, and George Floyd on social media” -Kwaning^10(p.1787)^

“By nature of our proximity to athletes, the orthopaedic subspecialty of sports medicine stands adjacent to this enormous potential…We’re privileged to enjoy a next-level degree of access. We should use our access to educate and empower. But too often, we miss the chance to engage, and we don’t have all the social impact that we might” -Owusu-Akyaw^26(p.672)^

“If you hear our voices, LISTEN to our pain. Do not just look at us, SEE our suffering. Do not just stand with us, SPEAK truth to power” -Mezu-Nbisi^24(p.4)^

“Now that we find ourselves at this place in history, we must purposely transition to becoming consciously competent.” -Vince^27(p.451)^

Authors largely discussed the phenomenon of racism at a distance, using impersonal nouns like “unconscious bias,” “discrimination,” and “impact” rather than referring to the specific people or institutions that perpetrated racism. However, whether articles were written by one or many individuals and whether or not white individuals were on the team, authors did write personally about themselves. They drew from their roles as Black physicians or physician trainees, positioning themselves as both medical experts and as experts–through personal experience–in bias, violence, and/or racist mechanisms like the minority tax and microaggressions. Yet, critically, they also aligned themselves with the broader, predominantly white medical community (i.e., their readers), shifting their stance throughout the articles.

While all these articles were selected in part because they had first-person singular “I” or first-person plural “we” pronouns, the ways in which authors used these pronouns varied. Some authors frequently referred to themselves in the first person (whether that was “I,” “we,” or a mix) while others primarily used third person references to other people, entities, or events. Authors also varied regarding when they presented first-person experiences. Some restricted this presentation to the beginning or the end of the article, while others wove personal pronouns fluidly throughout: talking about their own singular experiences with “I,” referencing a group of other Black individuals with “we,” and then naming themselves as part of a broader professional community.

When taking a first-person stance, authors described themselves in a variety of ways, including in relationship to their academic position like Doubeni’s naming of his Provost position (above); career stage like Carryl’s description of what she and others did “As Black medical students”^21(p.774)^ and medical specialty like Balzora’s naming of herself “As a Black female gastroenterologist.”^28(p.1).^ In this way, they made clear their location within the *profession* of medicine, often referring to their distance traveled to become physicians and leaders. This stance-taking acted, at least in part, to establish their credibility as part of the medical community and as valuable people, sometimes even referring to the quality of their family, as Mezu-Nbisi did (above).

These authors, like Griffin (above), also took stances as experts on *racism*. In telling these stories, some authors used strong affective language like Kwaning’s reference to trauma (above). Sharing their powerful personal “I” experiences and those of the Black community (or Black community of physicians) in America, these are places where authors demonstrated the authority of experience that allows them to critique the system.

Yet “we” did not always refer to the Black community–authors also used it to draw in a professional community that includes the reader like Owusu-Akyaw’s reference to all sports medicine doctors (above). In these cases, authors, despite their expressed stance as someone who has *experienced* racism, positioned themselves together with those white colleagues who may be part of *causing* those experiences. In fact, cases like Mezu-Dbisi’s use of second-person “you” (above) to call out the reader were rare. Instead, the reader is more often being called *in*, together with the author, as part of a“we” that can help address the racist transgressions. This calling in was often expressed through modals of obligation like Owusu-Akyaw’s use of “should” or through powerful rhetorical questions, implicating the reader in the solution.

### Intertextual Connections

“As Black physicians, we see the flawed healthcare system’s disproportionate and devastating effects on patients who look like us: we have first-hand accounts as patients ourselves, and we have traversed the experiences endured by our loved ones. Broken trust and fractured care contribute to disparate rates of morbidity and mortality in Black men and women with cardiovascular disease, stroke, and diabetes. ” -Unaka and Reynolds^29(p.572)^

“As a Black woman in medical school, I have had to cultivate ways to protect myself emotionally from the vicarious trauma triggered by witnessing the racially charged murders of Ahmaud Arbery, Breonna Taylor, and George Floyd on social media” -Kwaning^10(p.1787)^

How do we ease the fears of our mentees and ask them to trust our health and justice systems when curfews imposed by a pandemic put their lives in jeopardy (by increasing their chances of being pulled over by law enforcement officials)? Walter Scott died while running away from the police after five bullets punctured his body, including the fatal shot that traveled through his back to his heart. -Williams and Walker^30(p.192)^

“Unfortunately, medicine has a tradition of elevating and lauding physicians who have committed overtly racist acts (e.g., J. Marion Sims, Samuel A. Cartwright, Cornelius P. Rhoads).” -Shim^31(p.1795)^

“As Kēhaulani Kauanui reminds us, “Racism is a structure, not an event.” The tragic deaths of George Floyd and so many others are symptoms of deeply rooted racism that operates daily in our homes, neighborhoods—and our hospitals.” -Okaka and colleagues^32(p.202)^

“According to data from the Centers for Disease Control and Prevention [CDC], in 2018, the infant mortality rate for Black children was 10.8 per 1000 live births compared with a rate of 4.6 for nonHispanic white children.” -Adams, Davis, and Lechner^33(p.1)^

“In 2019, the Association of American Medical Colleges [AAMC] reported, “the growth of Black or African American [medical school] applicants, matriculants, and graduates lagged behind other groups.” -Flagg and Liu^34(p.241)^

Although none of the articles were empirical studies, no authors used *only* their personal experience in their pieces. Rather, to demonstrate both the value of what they were claiming and their *own* value as individuals, professionals, and witnesses to racism, authors aligned themselves through intertextual connections with patient care; social justice movements past and present; and scholarly and scientific structures.

To begin, several authors, like Unaka and Reynolds (above), referenced notions like patient care and safety, connecting their own experiences to those of the patients for whom they care. By taking a stance on racism’s harmful effects on patients, authors created value for their claims–this is not “just” about physicians and trainees, but about those who are at the center of medicine.

Authors also aligned themselves with social movements, including recent history that all their readers likely are aware of: COVID-19 and the disproportionate deaths of Black individuals (including naming some of those deaths like Susan Moore); police brutality and the deaths of Black individuals like Ahmaud Arbery, Breonna Taylor and George Floyd as in Kwaning (above); and the racist uprising on January 6th as Williams and Walker referenced (above). Authors also positioned themselves regarding past instances of racial violence: they critiqued the history over hundreds of years of slavery (Griffin referenced “shackles” in her title:

“Unshackled, but still Bound”^25^) and racism, including atrocities like Tuskegee and Marion Sims’ torture of Black women that Shim referenced (above). Finally, they aligned themselves with examples of Black agency and resistance, including activists like Kaepernick, Mandela, and Black Lives Matter; and Black scholars like Kendi, DuBois, Morrison and, as Okaka and colleagues did (above), Kauanui.

To further demonstrate validity of their claims, authors positioned themselves as scholars and scientists by drawing on the structures they hold dear. First, the majority of articles included cited references, including studies of healthcare disparities and inequity with indicators of how some academic field has covered the topic. As part of this alignment with the literature, some authors introduced academic constructs like “adult learning” or “servant leadership.” Authors, like Adams, Davis and Lechner (above), also frequently grounded their discussion in specific *statistics* and *facts*, often explicitly attributed to federal agencies like the CDC. Authors also referenced top professional entities like fields (e.g., pediatrics, PM&R), journals (e.g., *JAMA, Nature*), universities (e.g., Howard, Yale), well known hospitals (e.g., Massachusetts General, Deaconess) and other organizations, as Flagg and Liu did (above) when citing a trend noted by the AAMC.

### Promissory Space-Time Endings

“I hope for a more equitable and inclusive future for our field to which I am incredibly committed.” -Balzora^28(p.1)^

“Maybe the world will begin accepting that the confines in which we place each other for our own comfort were never meant to contain or define all that we are. Until the maybes become definite, I will continue to straddle the line between who I am and who I want to be. Maybe my perspective isn’t the only one that needs changing” -Langston^35(p.1979)^

“Let’s all inspire the change we want to see in our society. No more slow walking; we must pick up the pace. The time to do something is now!” -Ali^36(p.362)^

“With determination and discipline, we focus on the best ways to heal the wounds of racism and disparity inflicted upon all of us. We act in the midst of crisis because we are deeply devoted to and invested in our Black students and trainees. We reflect on those mentors and elders who showed the same devotion to us. As an act of gratitude and love, we steel ourselves to not falter in our dedication to mentoring. Together, we make a promise to each other that we never abandon this noble and worthy calling.” -Williams and Walker^30(p.193)^

“To my esteemed colleagues, I hope to leave you with just one lasting perspective. There is a loving family and a beautiful community coping with the loss of life, equality, and respect behind each data point of every disparities paper. My name is Dr. Carmen Black Parker, and I look forward to joining you in the literature.” -Parker^37(p.885)^

“As leaders of child mental health, we, as child psychiatrists, must do more. We, as health professionals, must do more. We must read more. We must learn more. We must speak more. The children are counting on us.” -Calhoun^38(p.145)^

“It is disappointing that research still sees Black and other students ‘treated as part of the homogeneous group’ [Black, Asian, and Minority Ethnic] BAME. We believe it is exactly this lack of delineation that hinders real change for Black medical students being implemented.” - Elewa-Ikpakwu and Ayoola^39(p.311)^

In all save one of the 29 articles, authors concluded by aligning themselves with the *future*, constructing a promissory or hopeful space-time with a nod to solutions, recommendations, or calls to action. These endings ranged from a hopeful call for change to institutions, programs, journals, departments, communities, leaders, or physicians, or even “the world” to a personal hope for change as Balzora did (above). While authors like Langston (above) tempered these calls with the difficulty of the current state of affairs, they still pointed to the promise of change. Sometimes authors, like Ali (above), included themselves in the latter call, using words such as “duty” even as they have narrated themselves as the targets of the racism that must be eliminated. This positioning as simultaneous agent and beneficiary of change was more explicit in some pieces, as with Williams and Walker (above), who restricted their message to their Black colleagues.

Turning to the beneficiaries of the hope or promise, many authors named Black or African-American groups like URM colleagues, Black high schoolers, Black people, or Black medical students. Others, like Parker (above), explicitly named themselves as benefitting from a hopeful future. But a number of articles shifted focus, with the last sentences of the article noting the change for us “all,” whether that be all people of all backgrounds, those in healthcare or, as Calhoun referenced, “the children.”

Yet one piece did not end this way; Elewa-Ikpakwu and Ayoola simply restated the problem, staying in that present troublesome space of racism rather than conforming to a hopeful narrative.

## Discussion

Because the discourses of medicine and medical publishing interpellate Black authors as Others and as potential “troublemakers,”^6^ they must carefully consider the stances they take, particularly when naming racism. The academic armor they put on must not only be able to defend them from attack, but to help them slip unseen through institutional bodies that are replete with mechanisms to eject them.^5^ Below we each reflect on our stances towards what we found, anchoring our piece with Author 1 while standing with her in resistance.

### Author 3: Language as Clue to Racial Strictures

As a white, heterosexual, cisgendered associate professor, I have the privilege of writing for publication without strategically shifting my stance between my professional and social communities. Black physicians and trainees already experience a fear of punishment or humiliation for raising concerns as racialized individuals in their work spaces^3^; our findings about how Black authors position themselves as medical professionals and experts in their personal narratives suggest that they may experience the same fear in publishing spaces. The emotional labor^8^ required to express the “authorial self”--”the extent to which a writer intrudes into a text and claims responsibility for its content”^40(p.1093)^ --likely discourages potential Black authors in medical education from sharing their stories.

Still, within academic medicine, published articles are critical for success and for promotion up the academic ladder. And writing establishes a particular kind of relationship that is perceived to be at the heart of academic identity: “textual production is at the core of negotiating the interactive relationships among the members of academic communities and claiming and constructing academic identities.”^16(p.83)^ In order to promote into places of power and influence, scholars must publish. I have always known that I am lucky that writing “comes easily” to me, but through this research I am realizing that it has little to do with some inherent writing ability or even learned writing skill. Instead, much of that ease derives out of the uncomplicated nature of my *stance:* when evaluating a construct like racism in my writing, I am not in danger of misalignment with white readers. I am not interpellated as Other. As I come to the end of this piece, however, I do feel an urge to use happiness as a way of “taking cover,” as “an obscurant”^6(p.83)^ of the trauma experienced by Black physicians and trainees, like my coauthor. But I am resisting that urge, ending instead with the words from the beginning of one of the articles by Alexis Griffin, her response after a long day at the hospital to the news of George Floyd’s death: “How much longer will we embrace this facade of the advancement of Black people? Could it be that we are still enslaved, but it just looks different? Will we ever truly be free, and how many people have to die before real change takes place?”

### Author 2: Intertextual connections as Academic Armor

As a white woman full professor and deputy editor-in-chief, through this study I saw for the first time how Black authors buttressed their personal experiences of racism with intertextual connections, many of which adhered to the traditions of academic discourse even in the absence of the structured IMRaD format. More specifically, as we discussed in our group the integration of statistics, federal agency reports, and professional association mandates made it feel like the authors were donning “academic armor”. This armor affords authors a protective layer of objectivity to surround their otherwise personal narrative, enabling them to adhere to the traditional expectations of a journal article.^14^ In 2001, Lerum described academic armor as “the physical and psychological means through which professional academics protect their expert positions or jurisdictions.”^41(p.470)^ Lerum describes that those clad in academic armor project emotional detachment and objectivity thus creating a “specific brand of knowledge”^41(p.475)^ which is prized in academic discourse. However, this approach discounts that academic writing is not just about conveying objective content, but that it is also about the representation of self.^40^

As an editor and professor, this realization made me wonder how I perpetuate authors’ sense that they must don academic armor. For example, often when editing, whether it be for a submitting author’s or my graduate student’s manuscript, I will add “(REF needed)” after a statement. Until now, I viewed this as a benign nudge, which I believed would ultimately help strengthen the work by aligning it with our field’s norms for journal articles. However, now I question this instinct. I hope that when faced with the temptation to add (REF needed), I will pause and ask, “Why do I feel that this author’s voice does not stand alone? Can/Should it?”. Additionally, I thought about the somewhat recent convention in health professions education encouraging authors to clearly state the “problem, gap, and hook”^42^ of their manuscript. While this can be a valuable heuristic it also conveys to authors that their personal experience may not be enough. Instead, to be published they must find an “important” problem supported by intertextual connections (e.g., patient care, social/historical movements). Beyond my individual editor role, this realization also made me consider the role of HPE journals. For example, author guidelines, which tend to be focused on research integrity and citation styles,^43^ say little about how to represent one’s personal experiences. Thus, it is unsurprising that authors continue to adhere to the rigid traditions of scholarly publishing highlighted in our journals and that reviewers and editors continue to expect it.

### Author 1: Reading Between the Lines

As Black woman medical student at a PWI, I see in these authors a similarity within myself as an author in the way they are aligning themselves with a hoped-for future rather than acknowledging the present. In all transparency, I approached this discussion the same way. I originally ended the discussion with the line “as such, there needs to be a space held for author’s stories that are held to a different standard,” a very structured sentiment grounded in hope for the future. It was only through debriefing with my mentors that they held me accountable in terms of not doing the same thing we were cautioning against. It is so ingrained in me to present information in this way as to not seem like the “angry Black woman” although I am not “happy and/or hopeful.” Even through rewrites of the draft it takes constant and intentional reflection as I write to make sure I am not following the status quo.

The vast majority (*n=28*) of the articles ended in the same fashion with some sort of positive ending. This brings up the question: are the authors actually hopeful about the future anti-racism efforts in medicine or is this also ingrained in them as authors or benignly suggested by mentors, reviewers, and editors in order to successfully publish? The way most of these authors don’t weave this promise of hope throughout and only use it briefly in the last paragraphs or sentences of their manuscript after detailing facts and experiences to portray the present, demonstrates the pull to end even a recounting of a traumatic experience in a positive way. As noted by Sarah Ahmed, marginalized individuals feel an institutional duty not to be negative or complain; they don’t want to come off as negative or complaining and want to be gentle on readers even in the midst of their trauma. Instead, I (we) must “swallow” these feelings because, “The more evidence you have that they are wrong, the more you are treated as being in the wrong, the harder they come down on you.”^5(p.48)^ This is how we are seen in society, when expressing the rawness of their experienced trauma, having to be conscious and careful of how it is perceived by the audience because if not received well, it will only exacerbate the trauma. Ahmed states, “We can think of narrative as a form of affective conversion. Through narrative, the promise of happiness is located as well as distributed. To make a simple point: some bodies more than others will bear the promise of happiness.”^6(p.45)^ The detrimental effects of not publishing in academia include not possessing as much academic currency as our white counterparts and is a driving force in conforming to academic standards to be published. Even writing now, I unconsciously don my academic armor as I write about the need to take it off and feel safe without it on because currently it does *not feel safe*.

### Author 1: Conclusions and Future Insights

Black physicians and trainees use various types of academic armor to amplify and validate their voices and experiences in academic writing. With this in mind, and exploring the standards mentioned above, we chose to open this paper with a vignette, in part to pay homage to the results we found, showing the importance of using one’s voice: that it is *enough*, it is *valid*, and it has an *impact*. This is a purposeful disruption of the traditional structure of academic work. The results of this paper bring several questions to the forefront… Are the discursive moves these authors make necessary when publishing about one’s experience? Are there more pieces from Black physicians and trainees that have been written but aren’t being published because they don’t have the appropriate armor? Does the culture of academic publishing make authors *feel unsafe*?

## Ethical Approval

Reported as not applicable

## Data

None reported

## Acknowledgement

Thank you to Joseph A. Costello for his assistance with manuscript preparation.

## Disclaimer

The views expressed in this article are those of the authors and do not necessarily reflect the official policy or position of the Uniformed Services University of the Health Sciences, the Department of Defense, or the U.S. Government.

## References

1. Althusser L. Ideology and Ideological State Apparatuses. Lenin and Philosophy and Other Essays. 1971. Translated by Brewster B. Available from: http://www.csun.edu/~snk1966/Lous%20Althusser%20Ideology%20and%20Ideological%20State%20Apparatuses.pdf

2. Johnson M, Konopasky A. Maintaining your voice as an underrepresented minority during the peer review process: A dialogue between author and mentor. Perspect Med Educ. 2022;11(3):144–145. doi:10.1007/s40037-022-00707-x.

3. Wyatt TR, Taylor TR, White D, Rockich-Winston N. “When No One Sees You as Black”: The Effect of Racial Violence on Black Trainees and Physicians. Acad Med. 2021;96(11S):S17–S22. doi:10.1097/ACM.0000000000004263

4. Wu J. Improving the Writing of Research Papers: IMRAD and Beyond. Landscape Ecology. 2011;26:1345–1349. doi:10.1007/s10980-011-9674-3.

5. Ahmed S. Complaint! Durham, NC: Duke University Press; 2021.

6. Ahmed S. The Promise of Happiness. Durham, NC: Duke University Press; 2010.

7. Torres MB, Salles A, Cochran A. Recognizing and Reacting to Microaggressions in Medicine and Surgery. JAMA Surg. 2019;154(9):868–872. doi:10.1001/jamasurg.2019.1648.

8. Osseo-Asare A, Balasuriya L, Huot SJ, et al. Minority Resident Physicians’ Views on the Role of Race/Ethnicity in Their Training Experiences in the Workplace. JAMA Netw Open. 2018;1(5):e182723. doi:10.1001/jamanetworkopen.2018.2723.

9. Zaidi Z, Partman IM, Whitehead CR, Kuper A, Wyatt TR. Contending with Our Racial Past in Medical Education: A Foucauldian Perspective. Teach Learn Med. 2021;33(4):453–462. doi:10.1080/10401334.2021.1945929.

10. Kwaning KM. Being Black in Medicine in the Midst of COVID-19 and Police Violence. Acad Med. 2020;95(12):1787–1788. doi:10.1097/ACM.0000000000003621.

11. Johnson M. Research as a Coping Mechanism for Racial Trauma: The Story of One Medical Student. Teach Learn Med. 2022;34(3):277–284. doi:10.1080/10401334.2021.1939033.

12. Sollaci LB, Pereira MG. The introduction, methods, results, and discussion (IMRAD) structure: a fifty-year survey. J Med Libr Assoc. 2004;92(3):364–367.

13. Froese FJ, Bader K. Surviving the desk-review. Asian Bus Manage. 2019;18:1–5. doi:10.1057/s41291-019-00060-8.

14. Gjesdal AM. The Influence of Genre Constraints on Author Representation in Medical Research Articles. The French Indefinite Pronoun On in IMRAD Research Articles. Discours [online]. 2013;12. doi:10.4000/discours.8770.

15. Fye WB. Medical authorship: traditions, trends, and tribulations. Ann Intern Med. 1990;113(4):317–325. doi:10.7326/0003-4819-113-4-317.

16. Flowerdew J, Wang SH. Identity in Academic Discourse. Annual Review of Applied Linguistics. 2015;35:81–99. doi:10.1017/S026719051400021X.

17. Du Bois JW. The stance triangle. In: Stancetaking in discourse: Subjectivity, evaluation, interaction. Englebretson R, eds. Amsterdam, NL: John Benjamins; 2007. (pg. 139–82).

18. Martin JR, Rose D. Working with discourse: Meaning beyond the clause. Sydney, AU: Bloomsbury Publishing; 2003.

19. Fairclough N. Analysing discourse: Textual analysis for social research. London: Routledge; 2003.

20. Bailey M, Mobley IA, Charles N, et al. Open Letter to Editors of Journal of the National Medical Association from the Black Feminist Health Science Studies Collective. J Natl Med Assoc. 2019;111(5):573–575. doi:10.1016/j.jnma.2019.04.003.

21. Carryl LM. Ways to Eradicate Systemic Racism in Health Care and Medical Education: A Letter to Medical Educators and Health Care Institutions. Acad Med. 2021;96(6):773–774. doi:10.1097/ACM.0000000000003986.

22. American Association of Medical Colleges. Diversity in Medicine: Facts and Figures 2019. Accessed: December 22, 2022. https://www.aamc.org/data-reports/workforce/report/diversity-medicine-facts-and-figures-2019.

23. Doubeni CA. Breaking Down the Web of Structural Racism in Medicine: Will JEDI Reign or Is It Mission Impossible? Mayo Clin Proc. 2021;96(6):1387–1389. doi:10.1016/j.mayocp.2021.04.017.

24. Mezu-Ndubuisi OJ. Unmasking Systemic Racism and Unconscious Bias in Medical Workplaces: A Call to Servant Leadership. J Am Heart Assoc. 2021;10(7):e018845. doi:10.1161/JAHA.120.018845.

25. Griffin A. Unshackled, but Still Bound: An Exploration of Racism in Medicine. Obstet Gynecol. 2020;136(5):1044–1046. doi:10.1097/AOG.0000000000004122.

26. Owusu-Akyaw K. The Forward Movement: Amplifying Black Voices on Race and Orthopaedics-It’s Time to Talk about Race in Sports Medicine. Clin Orthop Relat Res. 2021;479(4):671–673. doi:10.1097/CORR.0000000000001722.

27. Vince RA Jr. Eradicating Racial Injustice in Medicine-If Not Now, When? JAMA. 2020;324(5):451–452. doi:10.1001/jama.2020.12432.

28. Balzora S. When the minority tax is doubled: being Black and female in academic medicine. Nat Rev Gastroenterol Hepatol. 2021;18(1):1. doi:10.1038/s41575-020-00369-2.

29. Unaka NI, Reynolds KL. Truth in Tension: Reflections on Racism in Medicine. J Hosp Med. 2020;15(9):572–573. doi:10.12788/jhm.3492.

30. Williams DR, Walker VP. Curating Anger and Anguish Into Determination and Devotion: Black Women Faculty as Mentors in Medicine. Acad Pediatr. 2021;21(2):191–193. doi:10.1016/j.acap.2020.12.009.

31. Shim RS. Dismantling Structural Racism in Academic Medicine: A Skeptical Optimism. Acad Med. 2020;95(12):1793–1795. doi:10.1097/ACM.0000000000003726.

32. Okaka Y, AbdelHameid D, Olson RM, Kwarteng-Siaw M, Spanos N, Stone VE. Rallying Against Racism: Hospitals Join the Fight for Racial Justice. J Gen Intern Med. 2021;36(1):200–202. doi:10.1007/s11606-020-06291-2.

33. Adams SY, Davis TW, Lechner BE. Perspectives on Race and Medicine in the NICU. Pediatrics. 2021;147(3):e2020029025. doi:10.1542/peds.2020-029025.

34. Flagg CA, Liu MF. The Work Is Just Beginning-Racism in Medicine. Otolaryngol Clin North Am. 2021;54(1):239–245. doi:10.1016/j.otc.2020.09.018.

35. Langston AL. Dilemmas of Double Consciousness - On Being Black in Medicine. N Engl J Med. 2021;384(21):1978–1979. doi:10.1056/NEJMp2100211.

36. Ali S. COVID-19 Through The Eyes Of A Black Medical Student. Health Aff (Millwood). 2021;40(2):359–362. doi:10.1377/HLTHAFF.2020.01324.

37. Parker CB. Black in American Medicine: An Early-Career Psychiatrist’s Journey to Stand Against Disparities. Am J Geriatr Psychiatry. 2020;28(8):881–885. doi:10.1016/j.jagp.2020.04.006.

38. Calhoun A. Medical Education Must Start Teaching About Racism. Yale J Biol Med. 2021;94(1):143–146.

39. Elewa-Ikpakwu C, Ayoola G. Response to: Double jeopardy: Black and female in medicine. Clin Teach. 2021;18(3):311. doi:10.1111/tct.13306.

40. Hyland K. Authority and invisibility: Authorial identity in academic writing. Journal of pragmatics. 2002;34(8):1091–1112.

41. Lerum K. Subjects of Desire: Academic Armor, Intimate Ethnography, and the Production of Critical Knowledge. Qualitative Inquiry. 2001;7(4):466–483. doi: 10.1177/107780040100700405.

42. Lingard L. Joining a conversation: the problem/gap/hook heuristic. Perspect Med Educ. 2015;4(5):252–253. doi:10.1007/s40037-015-0211-y.

43. Malicki M, Jeroncić A, Aalbersberg IJ, Bouter L, Ter Riet G. Systematic review and meta-analyses of studies analysing instructions to authors from 1987 to 2017. Nature communications. 2021;12(1):1–4. doi:10.1038/s41467-021-26027-y.

