## Appendix A for "Putting on academic armor: How Black physicians and trainees take stances to make racism visible amidst publishing constraints"

PubMed

(("race"[Title] OR "racism"[Title] OR "anti-racism"[Title] OR "racist"[Title] OR "racial"[ti] OR "anti-racist"[Title] OR "Black"[Title] OR "African-American"[Title]) AND ("Medicine"[Title] OR "medical"[Title] OR "healthcare"[Title] OR "hospital*"[Title])) AND (comment[Filter] OR editorial[Filter] OR letter[Filter] OR personalnarrative[Filter] OR review[Filter]) AND (2018:2022[pdat])

PsycINFO

TI ("race" OR "racism" OR “racial” OR "anti-racism" OR "racist" OR "anti-racist" OR "Black" OR "African-American") AND TI ("Medicine" OR "medical” OR "healthcare" OR "hospital*")

*Limiters* - Document Type: Column/Opinion, Editorial, Letter, Review

*Date Limit:* 2018-2022

CINAHL

TI ("race" OR "racism" OR “racial” OR "anti-racism" OR "racist" OR "anti-racist" OR "Black" OR "African-American") AND TI ("Medicine" OR "medical” OR "healthcare" OR "hospital*")

*Limiters* - Publication Type: Anecdote, Commentary, Editorial, Interview, Letter, Response, Review

*Date Limit:* 2018-2022

Web of Science

TI=(("race" OR "racism" OR “racial” OR "anti-racism" OR "racist" OR "anti-racist" OR "Black" OR "African-American") AND ("Medicine" OR "medical” OR "healthcare" OR "hospital*"))

*Limiters* - Document Types: Letters or Editorial Materials or News Items or Notes or Reviews

*Date Limit*: 2018-2022
