## Appendix B for "Putting on academic armor: How Black physicians and trainees take stances to make racism visible amidst publishing constraints"

1. Nguemeni T. The Pictures on the Wall: Transitioning From a Historically Black College/University (HBCU) to an Ivy League Medical School. *Acad Med*. 2018;93(6):820. doi:10.1097/ACM.0000000000002191
2. Bailey M, Mobley IA, Charles N, et al. Open Letter to Editors of Journal of the National Medical Association from the Black Feminist Health Science Studies Collective. *J Natl Med Assoc*. 2019;111(5):573-575. doi:10.1016/j.jnma.2019.04.003
3. Parker CB. Black in American Medicine: An Early-Career Psychiatrist's Journey to Stand Against Disparities. *Am J Geriatr Psychiatry*. 2020;28(8):881-885. doi:10.1016/j.jagp.2020.04.006
4. Vince RA Jr. Eradicating Racial Injustice in Medicine-If Not Now, When? *JAMA*. 2020;324(5):451-452. doi:10.1001/jama.2020.12432
5. Griffin A. Unshackled, but Still Bound: An Exploration of Racism in Medicine. *Obstet Gynecol*. 2020;136(5):1044-1046. doi:10.1097/AOG.0000000000004122
6. Kwaning KM. Being Black in Medicine in the Midst of COVID-19 and Police Violence. Acad Med. 2020;95(12):1787-1788. doi:10.1097/ACM.0000000000003621
7. Owoseni AV. From Portraits to Role Models - Why We Need Black Physicians in Academic Medicine. N Engl J Med. 2020;383(23):2204-2205. doi:10.1056/NEJMp2027850
8. Okaka Y, AbdelHameid D, Olson RM, Kwarteng-Siaw M, Spanos N, Stone VE. Rallying Against Racism: Hospitals Join the Fight for Racial Justice. *J Gen Intern Med*. 2021;36(1):200-202. doi:10.1007/s11606-020-06291-2
9. Flagg CA, Liu MF. The Work Is Just Beginning-Racism in Medicine. *Otolaryngol Clin North Am*. 2021;54(1):239-245. doi:10.1016/j.otc.2020.09.018
10. Adams SY, Davis TW, Lechner BE. Perspectives on Race and Medicine in the NICU. *Pediatrics*. 2021;147(3):e2020029025. doi:10.1542/peds.2020-029025
11. Calhoun A. Medical Education Must Start Teaching About Racism. *Yale J Biol Med*. 2021;94(1):143-146.
12. Mezu-Ndubuisi OJ. Unmasking Systemic Racism and Unconscious Bias in Medical Workplaces: A Call to Servant Leadership. *J Am Heart Assoc*. 2021;10(7):e018845. doi:10.1161/JAHA.120.018845
13. Langston AL. Dilemmas of Double Consciousness - On Being Black in Medicine. *N Engl J Med*. 2021;384(21):1978-1979. doi:10.1056/NEJMp2100211
14. Elewa-Ikpakwu C, Ayoola G. Response to: Double jeopardy: Black and female in medicine. *Clin Teach*. 2021;18(3):311. doi:10.1111/tct.13306
15. Carryl LM. Ways to Eradicate Systemic Racism in Health Care and Medical Education: A Letter to Medical Educators and Health Care Institutions. *Acad Med*. 2021;96(6):773-774. doi:10.1097/ACM.0000000000003986
16. Doubeni CA. Breaking Down the Web of Structural Racism in Medicine: Will JEDI Reign or Is It Mission Impossible? *Mayo Clin Proc*. 2021;96(6):1387-1389. doi:10.1016/j.mayocp.2021.04.017
17. Ighodaro ET, Littlejohn EL, Akhetuamhen AI, Benson R. Giving voice to Black women in science and medicine. *Nat Med*. 2021;27(8):1316-1317. doi:10.1038/s41591-021-01438-y
18. Corbin TJ, Tabb LP, Rich JA, Kline JA. Commentary on Facing Structural Racism in Emergency Medicine. *Acad Emerg Med*. 2020;27(10):1067-1069. doi:10.1111/acem.14093
19. Williams DR, Walker VP. Curating Anger and Anguish Into Determination and Devotion: Black Women Faculty as Mentors in Medicine. *Acad Pediatr*. 2021;21(2):191-193. doi:10.1016/j.acap.2020.12.009
20. Charleston L 4th, Spears RC, Flippen C 2nd. Equity of African American Men in Headache in the United States: A Perspective From African American Headache Medicine Specialists (Part 1). *Headache*. 2020;60(10):2473-2485. doi:10.1111/head.14004
21. Norman J, Plummer CCJ 2nd. Moving From Asking to Action: We Must Do Better. *Am J Phys Med Rehabil*. 2021;100(3):e33-e34. doi:10.1097/PHM.0000000000001519
22. Owusu-Akyaw K. The Forward Movement: Amplifying Black Voices on Race and Orthopaedics-It's Time to Talk about Race in Sports Medicine. *Clin Orthop Relat Res*. 2021;479(4):671-673. doi:10.1097/CORR.0000000000001722
23. Unaka NI, Reynolds KL. Truth in Tension: Reflections on Racism in Medicine. *J Hosp Med*. 2020;15(9):572-573. doi:10.12788/jhm.3492
24. Shim RS. Dismantling Structural Racism in Academic Medicine: A Skeptical Optimism. *Acad Med*. 2020;95(12):1793-1795. doi:10.1097/ACM.0000000000003726
25. Golden SH. The perils of intersectionality: racial and sexual harassment in medicine. *J Clin Invest*. 2019;129(9):3465-3467. doi:10.1172/JCI130900
26. Ali S. COVID-19 Through The Eyes Of A Black Medical Student. *Health Aff (Millwood)*. 2021;40(2):359-362. doi:10.1377/HLTHAFF.2020.01324
27. Douglas A, Hendrix J. Black Medical Student Considerations in the Era of Virtual Interviews. *Ann Surg*. 2021;274(2):232-233. doi:10.1097/SLA.0000000000004946
28. Balzora S. When the minority tax is doubled: being Black and female in academic medicine. *Nat Rev Gastroenterol Hepatol*. 2021;18(1):1. doi:10.1038/s41575-020-00369-2
29. Vinekar K. Pathology of Racism - A Call to Desegregate Teaching Hospitals. *N Engl J Med*. 2021;385(13):e40. doi:10.1056/NEJMpv2113508
